# Chronicle of a Death Foretold: The Ecological Collapse of Sakumo Lagoon, Ghana

**DOI:** 10.64898/2026.07.14.738434

**Authors:** Margaret Fafa Awushie Akwetey, Emmanuel Lamptey, Sika Abrokwah, Denis Worlanyo Aheto, Paul Kojo Mensah, Isaac Okyere, Shehu Latunji Akintola, Daniel Pauly

**Affiliations:** Department of Fisheries and Aquatic Sciences, Africa Ocean Institute, University of Cape Coast, PMB Cape Coast, Ghana; Envaserv Research Consult, Sowutuom, Accra, Ghana; Centre for Coastal Management (CCM)/ Africa Centre of Excellence in Coastal Resilience (ACECoR), Africa Ocean Institute, University of Cape Coast, PMB Cape Coast, Ghana; Institute for Water Research, Rhodes University, Makhanda, 6140, South Africa; Fisheries Department, Lagos State University, Ojo, P.O. Box 001, LASU, Lagos, Nigeria; Sea Around Us, Institute for the Ocean and Fisheries, University of British Columbia, Vancouver, B.C., Canada, V6T 1Z4

**Keywords:** Benthic communities, Ecological collapse, Ramsar Site, Sakumo Lagoon, *Sarotherodon melanotheron*, Wetland

## Abstract

Sakumo Lagoon, a small (1 km^2^) semi-open coastal lagoon in Ghana, lies between the cities of Accra and Tema. The lagoon and its surrounding wetland were designated a Ramsar Site in 1992, mainly because it served as a refuge for 66 local and migratory bird species. Its ecology, and the biology of its major fish species, notably the blackchin tilapia (*Sarotherodon melanotheron*) were thoroughly studied in 1971, when the lagoon was a diverse, mainly brackish ecosystem supporting a traditionally and well-managed fishery. In 2016-2017, another study found the lagoon mostly covered by floating vegetation and plastic waste. Finally, in 2024, a visual survey established that the floating vegetation had been almost completely replaced by terrestrial plants, with only a few square meters of garbage-strewn water in front of a culvert connecting the lagoon to the open sea. Several lagoons along the coast of Ghana have been similarly lost to urban sprawl and its various forms of pollution, but Sakumo Lagoon is a Ramsar Site, and its imminent disappearance should not remain undocumented.

## 1. Introduction

There are uplifting stories of degraded aquatic ecosystems recovering their biodiversity and regaining a functioning ecosystem following near destruction. One beautiful example is narrated in ‘*The Death and Life of Monterey Bay: A Story of Revival’* by [1]. On the contrary, our story is not uplifting; rather, it describes the agony of a small West-African lagoon, from its already modified, but still ‘healthy’ state in 1971 to its heavily polluted and depauperate state in the late 2010s, then its demise in the 2020s, when the opportunity for an intervention to re-establish the functional structure of this past, vibrant ecosystem had vanished.

Lagoon ecosystems provide habitats, feeding, breeding, and nursery grounds for economically important fish species [2,3,4], and support productive fisheries [5]. They also function as roosting sites and stopovers for birds, especially migratory waterbird species [6]. In Ghana, as in other coastal communities globally, people living near coastal lagoons depend on them for shell- and fin-fish as major protein sources, mangroves for fuel wood, and recreation, among others. Additionally, lagoons provide other services such as protection of shorelines from storm surges and filtration of pollutants [7,8]. Despite their socio-economic and ecological importance, coastal lagoons in Ghana have come under severe threats in the last few decades [9,10,11], in part due to increasing human populations, which tend to concentrate in coastal areas. According to [12], despite lagoons having distinctive species of flora and fauna, as well as highly productive waters, they have been subjected to overexploitation and industrialization impacts. Consequently, many coastal lagoons in Ghana have been affected by various forms of pollution, leading to the loss of services and goods these ecosystems provide [13,14,15,16].

There are two coastal lagoons called ‘Sakumo’ in Ghana, both located in the Greater Accra Region. The bigger one is Sakumo I lagoon, which lies near the Densu delta. The one we will describe, being smaller, is often referred to as ‘Sakumo II,’ as did previous authors [17,18]. However, as we will not mention the bigger lagoon again, our ‘Sakumo Lagoon’ will refer to the smaller one (**Figure 1**). Sakumo Lagoon is in the Greater Accra region of Ghana, about 3 km west of Tema, the industrial hub of the country, and 1 km east of the Regional Maritime University in Accra, the country’s capital city (**Figure 2**).

**Figure 1.**
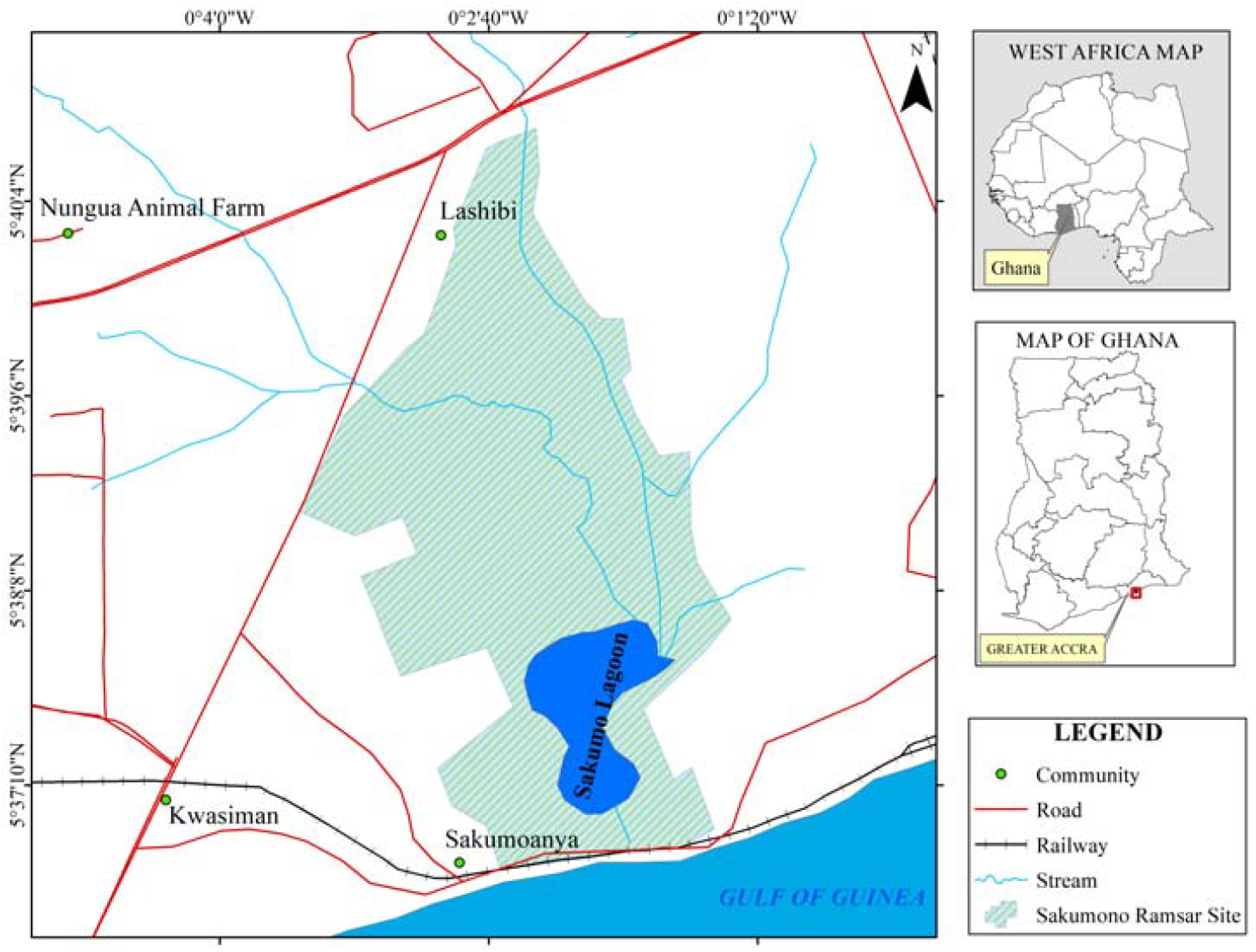
Geographical location of Sakumo Lagoon in the context of West Africa and Ghana.

**Figure 2.**
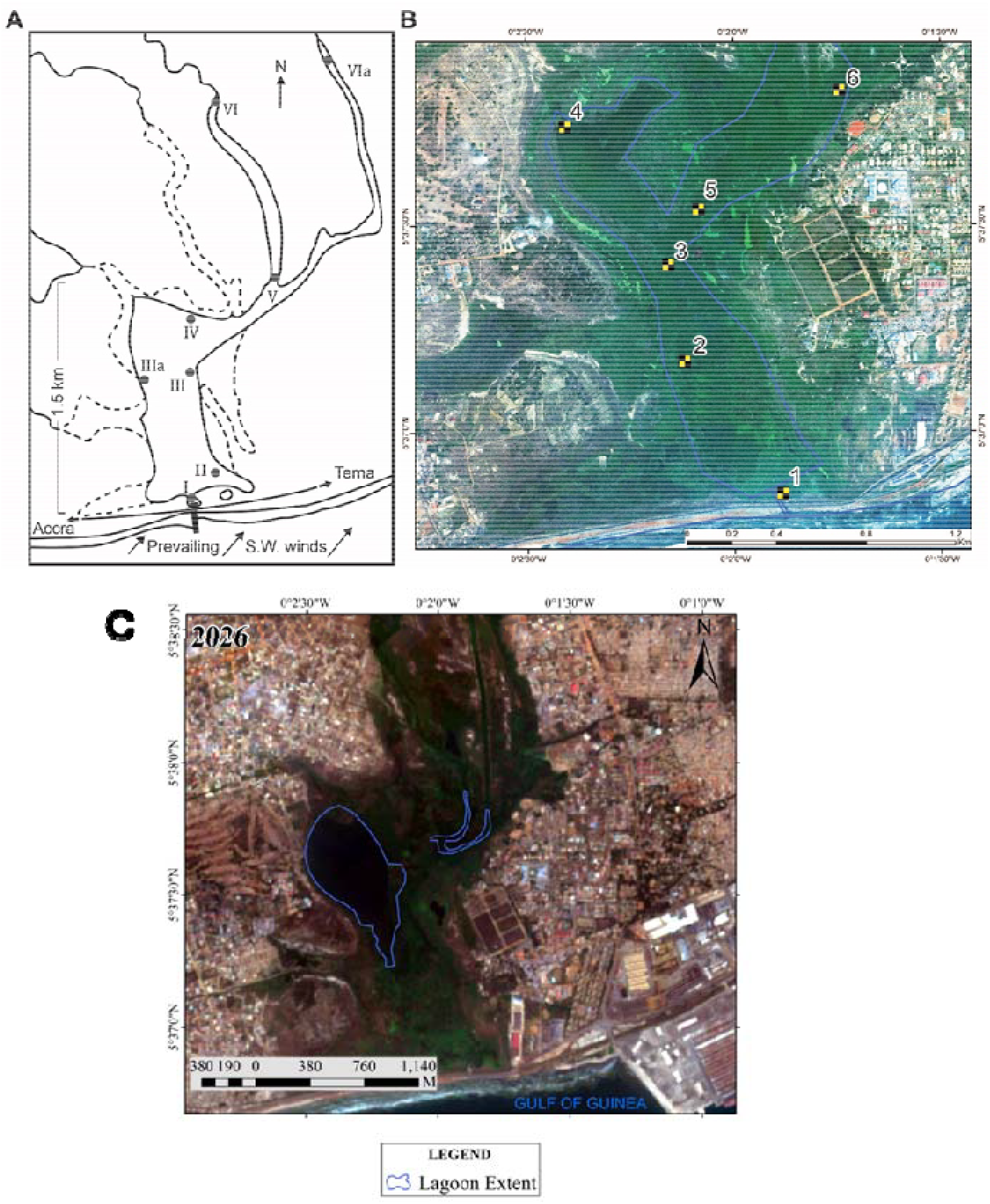
Sakumo Lagoon, Ghana. **A**: map of Sakumo Lagoon in the early 1950s, when its surface area was already reduced in extent compared to the second half of the 19^th^ century (see [19], figure 1). **B**: Satellite image of Sakumo Lagoon and the surrounding area in February 2016, with the 6 sampling stations in 2016/2017 showing human developments very close to the bank of the lagoon. **C**: Satellite image of the Sakumo Lagoon and its surroundings in 2026 (the outline of the open water in 2016 and 2026 is indicated in blue).

Sakumo Lagoon is part of the Sakumo wetland, which was internationally recognised among five coastal wetlands (Ramsar Sites) in 1992 under the Convention on Wetlands of International Importance in Ghana (**Figure 3**). The Ramsar Convention website [20] lists the services the Sakumo Lagoon provides as “cultural”; “provisioning”; “regulating” and “supporting” and mentions the existence of a ‘management plan’ for the lagoon. The designation of the Sakumo wetland as a Ramsar site was based on its status as a Key Biodiversity Area (KBA) of international significance, as over 80% of the waterbird species recorded there are palaearctic migrants [20]. Additionally, it provides support for migratory birds during critical life cycle stages (nesting and breeding) and in adverse conditions. In 1991, [21] reported that Sakumo Lagoon was the third most important site for shore birds along the coast of Ghana with 66 bird species, and an overall bird population of 32,500 individuals. Among them were six internationally important wader species, namely Spotted redshank, Greenshank, Curlew sandpiper, Little stint, Black-tailed godwit, and Black-winged stilt.

**Figure 3.**
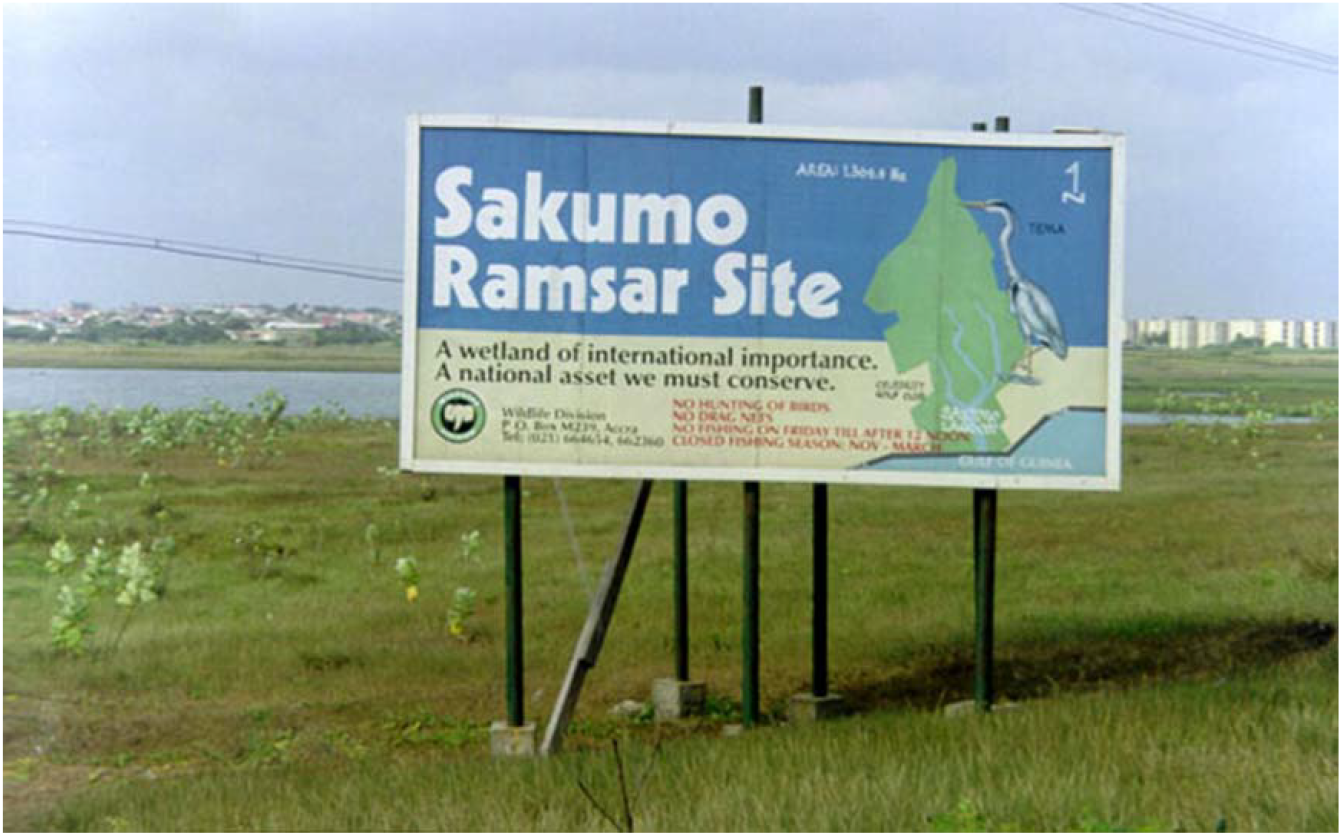
Signpost highlighting the Ramsar status of the Sakumo lagoon [Photo by Roger Pullin, 2008].

A review of the principal stipulations of the Ramsar Convention [22] concerning wetlands indicate in article 2 section 6 that ‘*Each Contracting Party shall consider its international responsibilities for the conservation, management and wise use of migratory stocks of waterfowl, both when designating entries for the List and when exercising its right to change entries in the List relating to wetlands within its territory’* implying a global interest when it comes to conservation and sustainable use of designated wetlands. In the early 1970s, Sakumo Lagoon was described as ‘semi-closed’ by [19], because it did not fit in the classification of [23], who distinguished, along the West African coast, two basic types of lagoons: the large ones, which are permanently open to the sea (e.g., Keta Lagoon; [24]) and a multitude of smaller ones which break the coastal sandbar and are open to the sea only during the rainy season, mainly from June to September. Sakumo Lagoon, while small, is connected to the sea via huge culvert pipes installed under the coastal road linking Accra to Tema, but they are now nearly blocked by sediments and refuse. The culvert enabled, throughout the years, a strong exchange of water and marine organisms between Sakumo Lagoon and the adjacent Gulf of Guinea, particularly at high tides. [19] described this exchange in some detail, as it was the reason for this small lagoon’s high biodiversity and productivity.

### Sakumo Lagoon in the late 1960s – early 1971

**Figure 2A** contrasts with the present extent of Sakumo Lagoon, illustrated in a recent satellite image (**Figure 2B**). Over 50 years ago, it was surrounded by isolated trees but lacked habitations along or even near its shores (**Figure 4A**).

**Figure 4.**
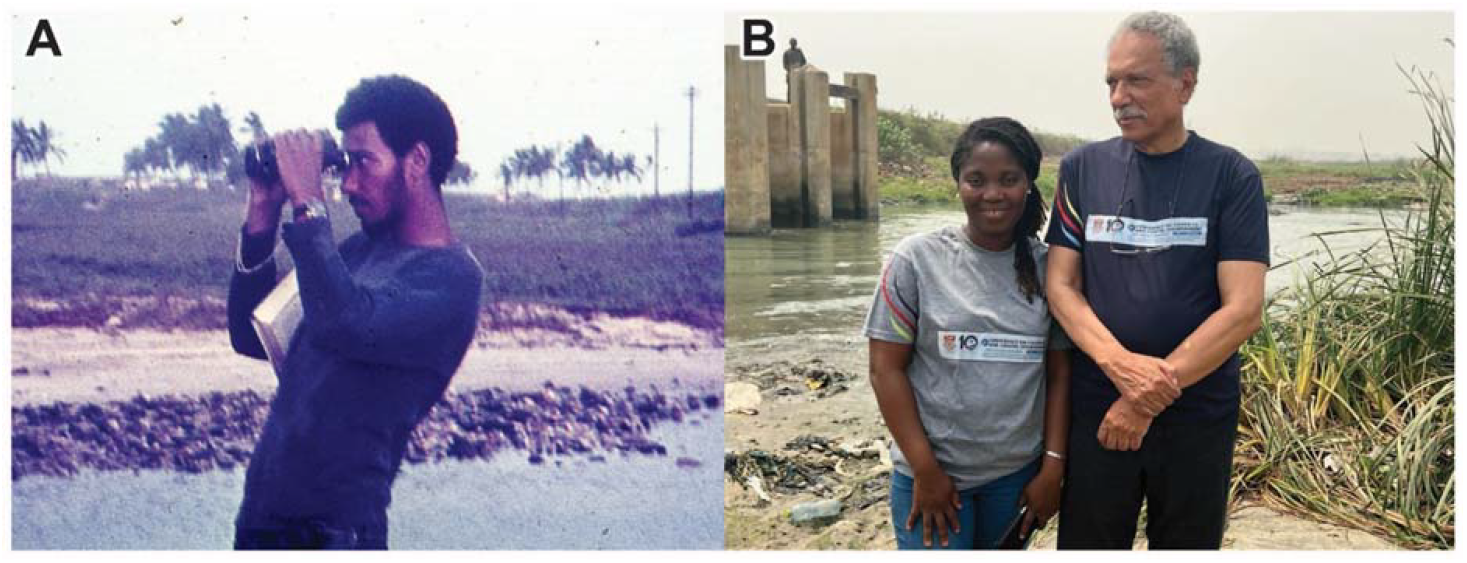
Two ages of investigations on Sakumo Lagoon. **A:** Daniel Pauly in 1971, when he studied the lagoon for his MSc thesis; **B:** the same, now 80 years old, with Margaret Fafa Awushie Akwetey, who led the re-investigation of the lagoon in 2016-2017; the water behind them is about 50% of the water surface of what is left of the lagoon as of February 2024 [Photo by Denis W. Aheto].

The central and northern part of the lagoon was the habitat of a thriving population of blackchin tilapia, *Sarotherodon melanotheron* (Rüppel 1852), which sustained a traditionally managed fishery (**Figure 5**). The local fishers exploiting Sakumo Lagoon set limits as to who could fish and with what gear [25]. The fishery was productive and apparently sustainable – except that by targeting larger adults, it caused the selection-induced reduction of adult sizes that occurs in many fisheries [26].

**Figure 5.**
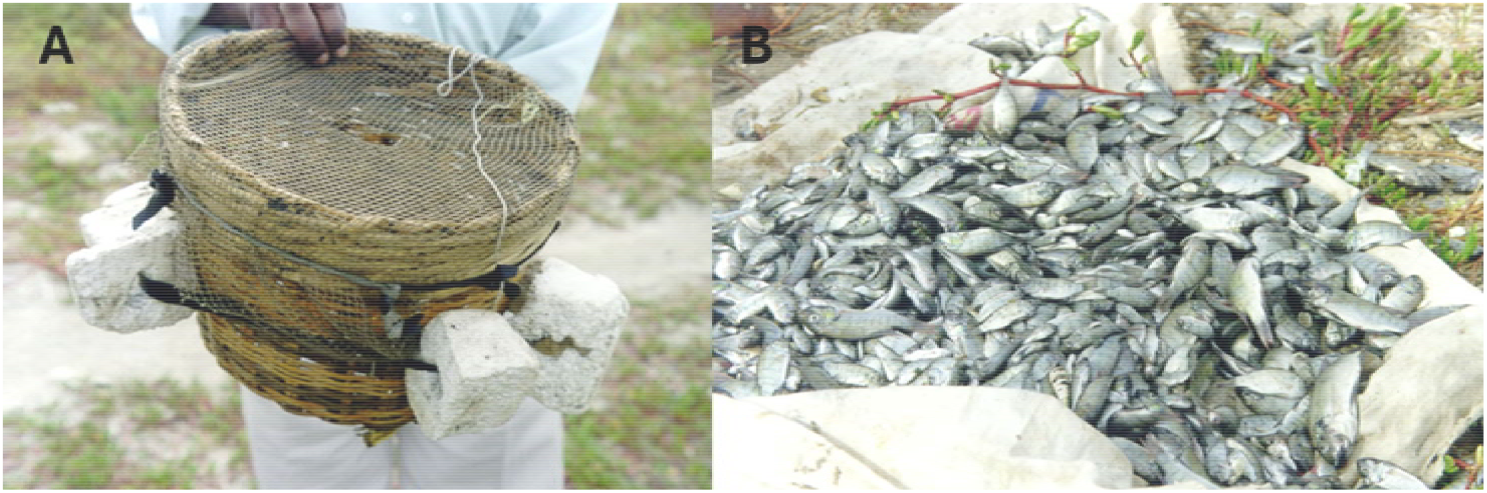
Blackchin tilapia *Sarotherodon melanotheron* caught from the Sakumo lagoon [Photo by Roger Pullin, 2008]

The southernmost part of the lagoon was strongly influenced by marine conditions due to the frequent water exchange through the culvert [19]. This led to an abundance of juvenile stages of commercially important marine fish species such as *Ethmalosa fimbriata, Mugil* spp., *Lutjanus* spp., and *Caranx* spp., which indicates the importance of this lagoon as a nursery ground for marine species. The molluscs in the lagoon were dominated by *Tympanotonus fuscatus* and *Crassostrea tulipa* (formally *Ostrea tulipa*), both of which were gathered and consumed locally [19].

The remarkable productivity of this fishery was based on the singular diet of the blackchin tilapia, which ingested large amounts of surficial mud at the bottom of the lagoon, from which their long gut extracted nourishment [25,27]. These fishes matured at about 10 cm, and rarely exceeded 15 cm total length, although they can reach a total length of over 30 cm in large Nigerian lagoons [28]. This seemingly stunted growth in the Sakumo Lagoon may be attributed to their parasites [29], the pressure of the intense fishery, which targeted large adults, as well as the large and stressful seasonal salinity and temperature fluctuations in Sakumo Lagoon, which far exceeded the fluctuations in open lagoon and in larger freshwater bodies, where the larger blackchin tilapia are found [28]. The last inference is supported by D. Pauly (pers. obs.) during a 1998 visit to Sakumo Lagoon, where fully mature specimens of 5-6 cm were observed, the diminutive descendants of the population studied in 1971 (see also [30], p. 194-195).

In 2002, [31] published an ecosystem model of Sakumo Lagoon constructed using the Ecopath with Ecosim (EwE) approach, based on the biodiversity data he had collected in 1971 and subsequently documented in 4 publications [19,25,27,29]. This model, which also had a spatial component, showed that a coherent account of the functioning of the Sakumo Lagoon ecosystem could be generated, i.e., that the main features of that ecosystem were understood.

Note, however, that Sakumo Lagoon, as studied in 1971, and described in the publications cited earlier, was not ‘pristine’; as a few tar balls were encountered along its banks, where crabs and other sand dwellers lived. While it was still functioning as an identifiable ecosystem, it had already been impacted by centuries of human exploitation and physical modifications. However, we remain unaware of these impacts, and the same will be true for the changes currently occurring, because the next generations will also experience a shift in their baseline [32]. This supports arguments that defining the ecological character of wetlands under Article 3, Section 2 of the Ramsar Convention on Wetlands, based on fixed reference conditions at the time of designation, may inadvertently restrict management strategies to already altered states and limit restoration possibilities [33]. A paleolimnological study of the Sakumo lagoon could eventually reveal insights into the ecosystem’s historical state, including its dynamics, biodiversity, timing, and direction of change over time, from decades to centuries. Such methods have yielded valuable understanding of sediment geochemistry and diatom distribution in East and South Africa [34,35], as well as the eutrophication and recovery of aquatic ecosystems in parts of the United States of America [36,37].

### The state of Sakumo Lagoon in the 21^st^Century

As mentioned above, Sakumo Lagoon is artificially connected to the sea by culvert pipes under the coastal road, with the direction of the flow depending on the tide [19,21,38], (see also background of **Figure 4B**) and it is fed freshwater by two streams, Mamahuma and Gbagbla Ankonu streams, which are heavily polluted with domestic wastes [18], and industrial wastes from companies including a Coca Cola bottling company [39]. In the mid-2020s, the banks of Sakumo Lagoon were used for vegetable farming, but the natural vegetation along its shores consisted mainly of *Sesuvium portulacastrum, Bothriochloa bladhii, Imperata cylindrica*, and *Typha domingensis* [21]. In 2016, the surface of the water was covered by water lettuce (*Pistia stratiotes*) for most of the year, which concealed the pollution stemming from domestic and industrial waste and effluents from a large catchment area within the Tema municipality (**Figure 6A**).

**Figure 6.**
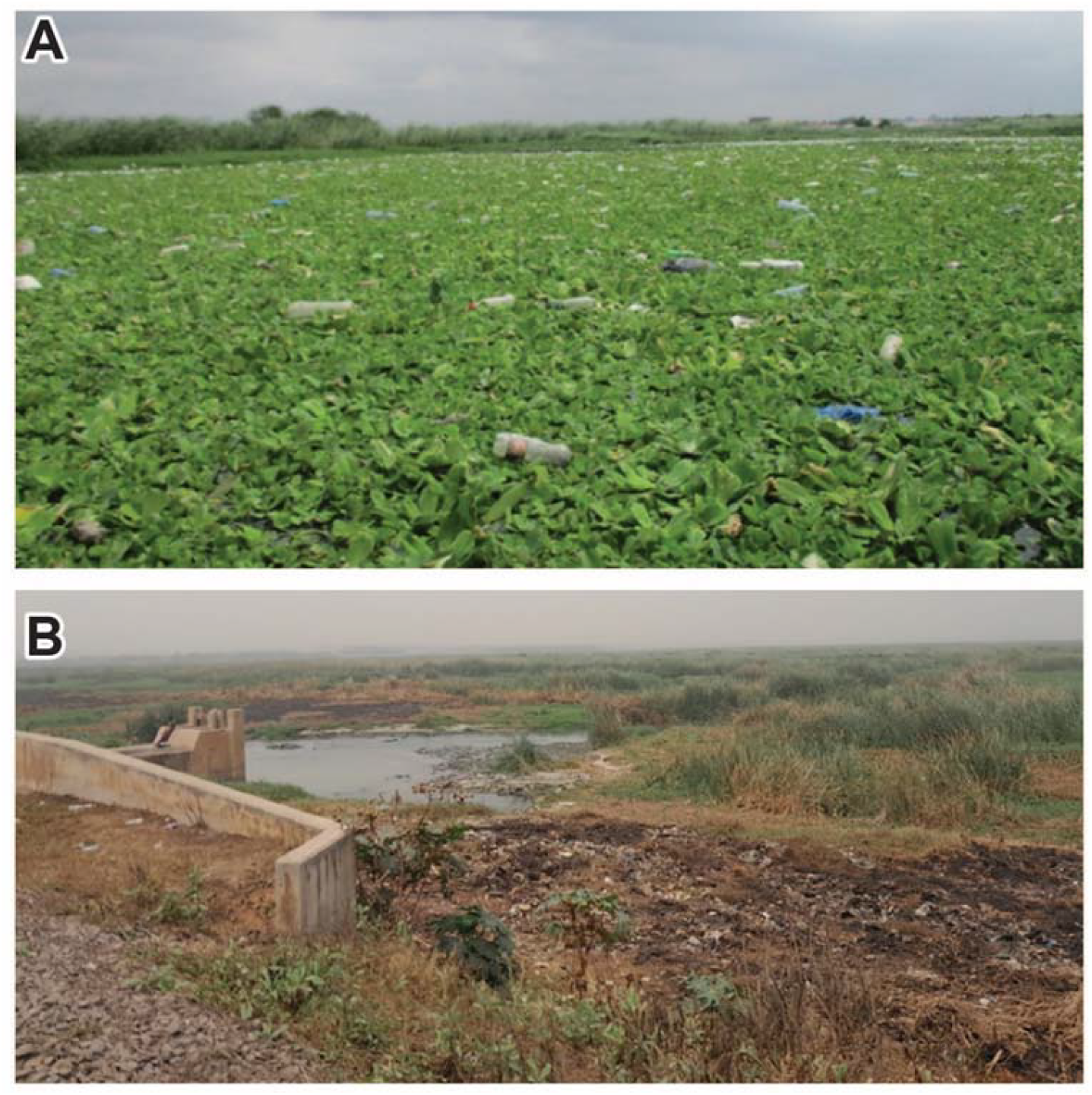
Two views of Sakumo Lagoon. **A:** ‘water’ surface in September 2016, covered by aquatic plants and plastic waste; **B:** the area that was the lagoon, in February 2024, is largely colonized by terrestrial plants [Photos by Margaret F. A. Akwetey]

Several studies have documented nutrient and heavy metal pollution in Sakumo Lagoon [15-18,40-44]. A 2008 study [40] attributed the lagoon’s hyper-eutrophic state to the impacts of human activities within its vicinity. [41] predicted that increasing nutrient loading to the lagoon would alter the structure and function of its biological communities, including benthic macroinvertebrates, thereby affecting the functioning of the lagoon’s ecosystem. Benthic macroinvertebrates are important components of the aquatic food web, acting as consumers of macrophytes and algae, and serving as food for fish. Their burrowing and feeding improve sediment aeration and stimulate nutrient regeneration, thus enhancing primary productivity [45-47]. However, only two records [19,48] of benthic macroinvertebrates are available for Sakumo Lagoon. In 1975, [19] reported the presence of polychaetes, bivalves, gastropods, and crustaceans, while a 1995 study [48] documented molluscs such as *Tympanotonus fuscatus, Turitella* sp., and *Macoma cumana* and three crab species, i.e., *Cardiosoma armati Callinectes amnicola* and *Uca tangeri*, all of which have now disappeared. Increasing pollution in the lagoon over the last three decades has altered its ecology, including the structure of the benthic macroinvertebrate community. This study aimed to highlight the ongoing degradation of the Sakumo Lagoon ecosystem using benthic macroinvertebrate assemblages as proxies for its overall ecological conditions.

## 2. Materials and Methods

The map of Sakumo Lagoon (5°37’24.27” N, 0°02’09.69” W) in Figure 2B shows the designated sampling stations. The depth of the lagoon ranged from 0.1 – 0.8 m during the sampling period. Sampling was conducted once at the end of each quarter from June 2016 to March 2017, with June and September corresponding to the wet season, whereas December and March correspond to the dry season.

Water parameters, including temperature (°C), dissolved oxygen (DO) concentration (mg/l), pH, salinity (‰), conductivity (µS/cm), and total dissolved solids (TDS) (ppm) were measured *in-situ* in triplicate at each station using a multi-parametric water quality checker (HI9829). The probe was positioned a few centimetres above the bottom. Turbidity (NTU) was measured with a turbidimeter (Oakton T-100). Water samples were collected from the same depth and sent to the laboratory for nitrate and phosphate analysis. Nitrate was determined using the cadmium reduction method, while the molybdovanadate method was used for phosphate determination. Sediment samples were also collected in triplicate using an Ekman grab and processed in the laboratory for organic matter (OM) and mean particle sizes (MPS) following methods described in [49]. Means and standard deviations of the physico-chemical parameters were computed, and a paired t-test was used to compare the parameters between wet and dry seasons.

Benthic macroinvertebrates were sampled with a 15 cm × 15 cm Ekman grab, washed through a 0.5 mm mesh sieve, preserved in a 10 % formalin solution, and stained with eosin to facilitate sorting in the laboratory. Three grab samples were taken at each sampling station. Fixed specimens were thoroughly rinsed with freshwater to remove the fixative, fine-grained particles, and debris. The sorted samples were examined under a dissecting microscope and identified to the lowest possible taxonomic level using identification guides [50-53]. The individuals of each species were recorded, and densities were calculated as the number of individuals per square meter. The density of species encountered in the two seasons was analysed for differences using a paired t-test.

## 3. Results

### 3.1. Environmental Parameters

In 2016-2017, polythene bags, plastic bottles, and cans were not only littering the surroundings of Sakumo Lagoon but were also floating on the surface of the lagoon, making it difficult to see its shoreline (**Figure 6A**). In 2024 and 2026, the water surface dried up and was largely taken over by terrestrial plants (**Figure 6B** and **Figure 2C** respectively).

Environmental parameters, including pH, dissolved oxygen (DO), salinity, nitrate, phosphate, and sediment mean particle size (MPS), showed significant differences between wet and dry seasons (**Table 1**). pH, DO, and phosphate measurements were significantly higher in the dry season, while salinity and nitrate were higher in the wet season. Sediment particle sizes were significantly larger (Coarse sand) in the wet season compared to the dry season, when sand of intermediate granularity was dominant.

**Table 1.** Differences in environmental parameters between wet and dry seasons.

| Physico-chemical parameters | Dry season (mean ± SD) | Wet season (mean ± SD) | t | p-value |
| --- | --- | --- | --- | --- |
| pH | 7.91 ± 0.66 | 7.53 ± 0.40 | -2.28 | 0.04* |
| DO (mg/L) | 4.53 ± 2.43 | 2.32 ± 0.52 | -3.04 | 0.01* |
| Temperature (°C) | 30.54 ± 1.24 | 29.50 ± 1.68 | -1.74 | 0.11 |
| Salinity (ppt) | 0.79 ± 0.19 | 3.25 ± 2.85 | 2.94 | 0.01* |
| Conductivity (μS/cm) | 1693.44 ± 429.26 | 1650.39 ± 257.58 | -0.41 | 0.69 |
| TDS (mg/L) | 804.64 ± 183.71 | 822.69 ± 128.71 | 0.41 | 0.69 |
| Turbidity (NTU) | 53.84 ± 38.49 | 247.61 ± 298.42 | 2.25 | 0.05 |
| Nitrate (mg/L) | 2.09 ± 1.62 | 14.02 ± 10.09 | 3.93 | 0.00* |
| Phosphate (mg/L) | 17.29 ± 15.36 | 1.84 ± 1.83 | -3.86 | 0.00* |
| Organic Matter (OM) | 6.57 ± 3.96 | 9.93 ± 5.38 | 1.78 | 0.10 |
| MPS (mm) | 0.40 ± 0.17<sup>a</sup> | 0.78 ± 0.19<sup>b</sup> | 6.26 | 0.00* |

df = 11, \* Indicates significant values ( $p < 0.05$ ), 'a' denotes Medium sand, 'b' denotes Coarse sand.

### 3.2. Benthic macroinvertebrates

Five (5) benthic macroinvertebrate species, consisting of 2 polychaetes, 1 oligochaete and 2 insect larvae were encountered in the 2016/2017 study compared to 20 species in 1971 (**Table 2**; see also **Tables 3** and **4**). Gastropods, bivalves and crustaceans, which were present in the previous study, were absent from this study. Abundance and mean density (± standard deviation) of benthic macroinvertebrates are in Tables 3 and 4, respectively. The large standard deviations observed across species indicate high variability in macroinvertebrate densities within the lagoon due to patchy distribution across stations.

**Table 2.** Species list of benthic macroinvertebrates^1^ in Sakumo Lagoon in 1971 and 2016/2017.

| Major group/species | 1971 | 2016/17 | Major group/species | 1971 | 2016/17 |
| --- | --- | --- | --- | --- | --- |
| **Polychaeta** |  |  | **Gastropoda (continued)** |  |  |
| *Syllis hyaline* | + | - | *Aplysia dactylomela* | + | - |
| *Notomastus lineatus* | + | - | **Bivalvia** |  |  |
| *Nephtys pyrifanis* | + | - | *Arca senilis* | * | - |
| *Aricidea* sp. | - | + | *Crassostrea tulipa* | + | - |
| *Capitella capitata* | - | + | **Crustacea** |  |  |
| **Oligochaeta** |  |  | *Balanus* spp. | * | - |
| *Limnodrilus hoffmeisteri* | - | + | *Siriella* spp. | + | - |
| **Insecta** |  |  | *Brandidirella megua* | + | - |
| *Notonecta* sp. | - | + | *Ligia exotica* | + | - |
| *Chironomus* sp. | - | + | *Penaeus duorarum* | + | - |
| **Gastropoda** |  |  | *Penaeus kerathurus* | + | - |
| *Tympanotonus fuscatus* | + | - | *Clibanarius africanus* | + | - |
| *Thais* sp. | + | - | *Callinectes latimanus* | + | - |
| *Glycymeris* sp. | + | - | *Uca tangeri* | + | - |
| *Pugilina morio* | + | - | *Cardisoma armatum* | + | - |
| *Peringia ulvae* | + | - | *Gionopsis cruentata* | + | - |

<sup>1</sup> Except for the insects, all species names in this table were verified against the valid names in the World Register of Marine Species [54] and entered into the records for Ghana in SeaLifeBase [55]; + indicates presence and - indicates absence, while \* indicates that only shells were found.

**Table 3.** Abundance of benthic macroinvertebrates in Sakumo Lagoon in 2016/2017.

| Species | Wet season | Dry season |
| --- | --- | --- |
| *Limnodrilus hoffmeisteri* | 486 | 1287 |
| *Capitella capitata* | 4 | 0 |
| *Aricidea* sp. | 1 | 0 |
| *Chironomus* sp. | 1094 | 802 |
| *Notonecta* sp. | 45 | 17 |

**Table 4.** Density of benthic macroinvertebrates in Sakumo Lagoon in 2016/2017.

| Species | Wet season (ind·m<sup>-2</sup>) | Dry season (ind·m<sup>-2</sup>) | t | p-value |
| --- | --- | --- | --- | --- |
| *Limnodrilus hoffmeisteri* | 599 ± 1726 | 1587 ± 2712 | -1.02 | 0.33 |
| *Capitella capitata* | 4.93 ± 11.5 | 0 | 1.48 | 0.17 |
| *Aricidea* sp. | 1.23 ± 4.27 | 0 | 1.0 | 0.34 |
| *Chironomus* sp. | 1349 ± 2430 | 989.1 ± 2038 | 0.47 | 0.65 |
| *Notonecta* sp. | 55.5 ± 161 | 21.0 ± 56 | 0.68 | 0.51 |

Values are presented as mean ± standard deviation (SD); density expressed as number of individuals per square meter (ind·m<sup>-2</sup>)

## 4. Discussion

A comparison with historical data indicates a significant ecological collapse of Sakumo Lagoon, evidenced by reduced macroinvertebrate diversity and a shift towards pollution-tolerant species. Pollution and loss of lagoons and other wetlands, which were once a habitat for many aquatic organisms, have been attributed to the spread and emergence of urbanization within the country [56]. Since 1982, Sakumo Lagoon has been listed among Ghana’s most polluted lagoons [10,14-16,40]. In 1991, [44] reported heavy littering of the immediate surroundings of the Sakumo lagoon with plastics, empty cans, and bottles, among others. Besides farming activities in the area, parts of the lagoon were transformed into a huge refuse dump, gradually killing aquatic life, and causing a shift in the benthic community structure in Sakumo Lagoon between 1971 and 2016/2017. Also, there was a massive decrease in species richness in this study for the same reason, confirming the prediction of [41] that the increasing pollution in Sakumo Lagoon, particularly nutrient loading, would impact the structure and function of biological communities of the lagoon.

The high conductivity of the water, coupled with high nitrate and phosphate concentrations recorded in this study as well as in previous work between the 1980s and 2010s [15-17,40-43] are also indicative of the deterioration in the ecological state of the lagoon. Conductivity above 1000 µS/cm has been linked to saline conditions and/or heavy impact by industry [57-59]. However, salinity values in this study were generally low (< 3 ‰), supporting the argument that high conductivity in Sakumo Lagoon is more likely due to pollution than salinity, which, it should be recalled, could reach high values [19]. High concentrations of nitrate and phosphate are likely leaching from municipal and industrial wastes, as well as farms along the banks of the lagoon. This is confirmed by [6], who noted that nutrient enrichment can be derived from different human activities and sources such as input of sewage, leaching of fertilizer, runoff from agricultural fields, etc., and could be more intense in shallow lagoons with little water renewal due to intermittent or no connection with the sea. The effects of excessive nutrients in Sakumo Lagoon were evident in 2016/2017 in the characteristic dark colour of the water and the bloom of invasive weeds on the surface of the water body.

Environmental conditions such as pollution, industrialization, urbanization, and eutrophication influence the distribution of species in the benthic community [60]. Benthic macroinvertebrate communities are critical in lagoons for maintaining ecosystem functioning through their roles in nutrient cycling, sediment structuring, energy transfer in food chains, etc. [6]. In the 2016/2017 study, two species (*Aricidea* sp. and *Capitella capitata*) present in the wet season disappeared in the dry season, leading to lower species richness in that season. The presence of these two species in the wet season could simply be an accidental occurrence, possibly from relics of a previous community. *Limnodrilus hoffmeisteri* and *Chironomus* sp. dominated the fauna in both dry and wet seasons. These species have been described as pollution-tolerant by many authors [50,52], hence their dominance in the lagoon. Large numbers of chironomid larvae are documented to thrive in polluted and eutrophic ecosystems due to their resilient and rapid colonizing capabilities [60-62]. Similarly, *L. hoffmeisteri* (Family Tubificidae) is tolerant to a variety of stress and serves as an indicator of eutrophic conditions [63]. Densities above 1000 ind. m^-2^, as observed in this study, are indicative of moderately enriched environments [64]. The poor benthic community structure in the lagoon supports the claim based on nutrient and conductivity values that Sakumo Lagoon was in a poor ecological state. A poorly structured benthic macroinvertebrate community has implications such as reduced nutrient cycling, decline in bioturbation and sediment restructuring, lower food availability for higher trophic levels, and subsequent decline in fisheries productivity [6,65,66].

In recent times, the deteriorating state of Sakumo Lagoon has been reported in both print and electronic news outlets in Ghana. For instance, [67] covered a story on these developments, highlighting the observed decline in the populations of migratory bird species that visit the lagoon seasonally. Similarly, [68] reported on the sad state of Sakumo Lagoon as a Ramsar Site, lamenting that a large portion of the lagoon was being reclaimed for estate developments. The reports also asserted that these developments have led to the decline of populations of migratory bird species that visit the lagoon seasonally. A similar concern was expressed by [69], emphasizing that the same estate development activities in the wetland are interfering with its ecological service as a flood control ecosystem, which is of great concern in the face of climate change with increasing sea levels. This situation confirmed earlier reports by [70], that hundreds of nearby houses were submerged due to continuous rainfall attributed to climate change. [71] further noted in their publication that besides the activities of encroachers and estate developers, parts of the lagoon are being used as refuse dumps, which is gradually harming aquatic life in the wetland.

Note that [38] in a report documenting the management plan for Sakumo Ramsar Site, following its designation as a Ramsar Site, outlined the values of this lagoon and the entire area. Sakumo Lagoon was a habitat for over 60 bird species, including 6 internationally important wader species, 11 distinct vegetation types, and 13 finfish species, some of which were of marine origin. The lagoon provided provisioning services including food (fish), fuelwood (mangroves), medicine (neem trees), etc. The banks of the lagoon were also used for vegetable farming. In terms of regulatory services, the vegetation within the area provided climate control services, the birds contributed to pollination, while the flood plains of the lagoon absorbed water that could have ended up in homes during the wet season. Finally, Sakumo Lagoon held cultural and religious significance, which has diminished over time due to increasing urbanization. The lagoon served as a home to a local deity while providing opportunities for ecotourism due to the migratory birds (see also [28]).

An update on the Ramsar Convention website [20] indicates that the Sakumo Ramsar Site, as an urban wetland, faces threats from population growth, urbanization, pollution, and development activities such as agriculture and recreation. Further, it is alluded that the small size of the lagoon and its location between two industrial cities made it vulnerable to the pressure of anthropogenic activities within the catchment [38]. The degradation of Sakumo Lagoon and other Ramsar sites can be linked to Article 2, Section 3 of the Ramsar Convention, which states that ‘*The inclusion of a wetland in the List does not prejudice the exclusive sovereign rights of the Contracting Party in whose territory the wetland is situated’* [22]. This suggests that the conservation of Ramsar sites largely depends on domestic governance and enforcement. When countries fail to protect these sites, pollution often results, and there are no significant penalties or deterrents. The success of protected areas in achieving conservation goals depends not only on their designation but also on governance frameworks, management capabilities, and the integration of ecosystem service considerations into decision-making [72]. A 2022 study [73] corroborates this, finding that protected area designation does not guarantee long-term ecological persistence under urbanization and climate stress. Systematic assessments of conservation effectiveness frameworks have also highlighted similar concerns about the limited ability of protected area designations to prevent biodiversity loss (74). The continued degradation of Sakumo Lagoon underscores broader concerns that Ramsar designation alone is insufficient without effective management and enforcement [75]. In Ghana, decision makers have failed to take action to safeguard these vital ecosystems. This failure could be attributed to the lack of relevant non-governmental organizations (NGOs) focused on wetland protection. NGOs provide key information for governments on issues of human interest, such as environmental degradation, while also providing a medium for the public to be involved in wetland conservation through activities such as awareness creation, education, volunteerism, etc. [76]. The absence of NGOs that could have acted as mediators between local communities and local authorities was also evident when the media issued several reports on the dying of Korle lagoon [77,78] and Chemu lagoon [79-81]. Evidence from other Ramsar sites indicates that community-driven conservation efforts led by NGOs can greatly enhance wetland management through stakeholder engagements, long-term monitoring, and institutional collaboration [82].

The Sakumo Ramsar site has experienced significant ecological decline since it was first designated. A review of the main provisions of the Ramsar Convention [22], Article 3, Section 2, indicates that ‘*Each Contracting Party shall arrange to be informed at the earliest possible time if the ecological character of any wetland in its territory and included in the List has changed, is changing or is likely to change as the result of technological developments, pollution or other human interference. Information on such changes shall be passed without delay to the organization or government responsible for the continuing bureau duties specified in Article 8’*. The Convention’s concept of “ecological character” highlights that many wetlands were already degraded when they were listed. Evidence [10,14–16,40,42,44] indicates that the Sakumo Lagoon was already under significant environmental stress prior to its designation in 1992 as an important habitat for migratory waterbirds, underscoring that its Ramsar status was conferred against a backdrop of pre-existing ecological degradation [33].

[83] explained that a contracting party is free to delist or restrict the boundaries of a Ramsar site, after following proper procedures to inform the Ramsar Bureau of the changes. The authors, however, noted that at the time of their report, no country had ever deleted a wetland from the list. Many contracting parties or states have listed Ramsar sites with changing ecological conditions due to human-induced threats on the Montreux Record. Established in 1990, the Montreux Record serves as a vital Ramsar mechanism to identify sites requiring urgent conservation actions and to guide resource allocation through existing financial channels. A site can be added to this record either at the request of the Party (i.e., the country) where the wetland is located or initiated by partner organizations, including international or national NGOs or other interested entities. However, such addition always requires approval by the concerned party. Ghana should consider listing the Sakumo Ramsar site on the Montreux Record. It is also noteworthy that other Ramsar sites in Ghana, such as Songhor, Keta, and Muni, are quietly experiencing severe degradation due to overexploitation and urbanization [12, 84-87].

A quick search on the Ramsar website [88] revealed 46 Ramsar sites on the Montreux Record; the majority (21) are in Europe, 10 in Asia, 8 in Africa, 6 in Latin America and the Caribbean, and one in North America. Among the threats common to these wetlands are natural system modifications, pollution, water regulation, biological resource use, agriculture and aquaculture, human settlements, etc. Notably, no Ramsar site in West Africa is on the Montreux Record. Two sites each were listed for Egypt (Lake Burullus and Lake Bardawil), South Africa (Blesbokspruit and Orange River Mouth), and the Democratic Republic of Congo (Parc national des Virunga and Parc national des mangroves), while one site was listed for Tunisia (Ichkeul) and Uganda (Lake George). Considering the ongoing environmental degradation in many West African wetlands, the fact that no wetland in the region is listed on the Montreux Record is questionable.

Regarding the Sakumo area, a section of the public suggested dredging the lagoon to help restore the flood control function of the wetland, as noted in [71]. Furthermore, numerous calls were made to restore Sakumo Lagoon to its earlier state, as echoed by [89-91], after a heavy downpour in June 2016 exposed the vulnerability of the lagoon’s adjoining communities. However, it does not appear that the competent authorities are committed to calls [92,93], which, if nothing is done, will mean the death of Sakumo Lagoon, as in the case of the Korle, Chemu, and Kpeshie lagoons [42,94,95].

## 5. Conclusion

The Sakumo Lagoon has shifted from a vibrant brackish water ecosystem to a polluted and altered freshwater eutrophic system, and then to a terrestrial habitat, exposing a gap between Ramsar recognition and the actual situation. While the Ramsar Convention offers a framework for the conservation and ‘wise-use’ of wetlands, the primary responsibility for their protection rests on contracting parties. It is also evident that West African countries do not utilize the Montreux Record and the Sakumo Lagoon urgently needs to be added to this record. Additionally, efforts should focus on restoring the lagoon’s ecological condition. The imminent loss of Sakumo Lagoon signals a disappearance of a Ramsar Site, resulting in biodiversity loss, economic decline, and livelihood challenges. This situation serves as a wake-up call for national authorities and international agencies, especially the Ramsar Convention, to intensify efforts to monitor and safeguard wetlands and their biodiversity, particularly those designated as Ramsar sites. The Ramsar Convention should consider revising its policies to delist contracting parties that fail to protect their Ramsar sites, thus remaining true to the objectives of the convention.

## Author Contributions

Margaret Fafa Awushie Akwetey – Conceptualization, Methodology of 2016/2017 studies, Investigation of 2016/2017, Formal analysis, Fund acquisition, Writing – original draft, Writing – review and editing. Emmanuel Lamptey – Conceptualization, Supervision, Writing – review and editing. Sika Abrokwah – Conceptualization, Writing – original draft. Denis Worlanyo Aheto – Fund acquisition, Formal analysis, Writing – review and editing. Paul Kojo Mensah – Formal analysis, Writing – review and editing. Isaac Okyere – Formal analysis, Writing – review and editing. Shehu Latunji Akintola – Formal analysis, Writing – review and editing. Daniel Pauly – Conceptualization, Methodology of 1971 studies, Investigation of 1971 studies, Writing – original draft, Writing – review and editing. All authors have read and agreed to the published version of the manuscript.

## Data Availability Statement

The original contributions presented in this study are included in the article. Further inquiries can be directed to the corresponding author.

## Acknowledgments

The authors are grateful to the late Professor Emeritus Kobina Yankson for providing guidance during the conceptualisation and data collection of 2016-2017, Elaine Chu for assembling our photos into publishable figures, Martin Efemor Paku for the satellite images and Roger Pullin for photos. Daniel Pauly uses this opportunity to thank Professor G. Hempel for the support he provided in 1971.

## References

1. Palumbi, S.R.; Sotka, C. The Death and Life of Monterey Bay: A Story of Revival; Island Press: Washington, DC, USA, 2011.

2. Simboura, N.; Maragou, P.; Paximadis, G.; Kapiris, K.; Papadopoulos, V.P.; Sakellariou, D.; Pavlidou, A.; Hatzianestis, I.; Salomidi, M.; Arvanitidis, C.; Panayotidis, P. Greece. In World Seas: An Environmental Evaluation Volume I: Europe, the Americas and West Africa, 2nd ed.; Sheppard, C., Ed.; Elsevier: 2018; pp. 227–260. 10.1016/B978-0-12-805068-2.00012-7

3. Vollum, D.; Coasts, A.; Vollum, K.S.D.; Koomson, D.; Raha, D. Coastal lagoons of West Africa: A scoping study of environmental status and management challenges. Anthropocene Coasts 2024. 10.1007/s44218-024-00039-9

4. Clark, J.R. Integrated management of coastal zones. FAO Fish. Tech. Pap. 1992, 327. Available online: http://www.fao.org/docrep/003/t0708e/t0708e03.htm (accessed on 24 May 2021).

5. Pauly, D.; Yañez-Arancibia, A. Fisheries in coastal lagoons. In Coastal Lagoon Processes; Kjerfve, B., Ed.; Elsevier Science Publishers: Amsterdam, The Netherlands, 1994; pp. 377–399.

6. Rodrigues-Filho, J.L.; Macêdo, R.L.; Sarmento, H.; Pimenta, V.R.A.; Alonso, C.; Teixeira, C.R.; Pagliosa, P.R.; Netto, S.A.; Santos, N.C.L.; Daura-Jorge, F.G.; Rocha, O.; Horta, P.; Branco, J.O.; Sartor, R.; Muller, J.; Cionek, V.M. From ecological functions to ecosystem services: Linking coastal lagoons biodiversity with human well-being. Hydrobiologia 2023, 850, 2611–2653. 10.1007/s10750-023-05171-0

7. El Mahrad, B.; Newton, A.; Murray, N. Coastal lagoons: Important ecosystems. Frontiers for Young Minds 2022, 10. 10.3389/frym.2022.637578

8. Massarelli, C.; Campanale, C.; Uricchio, V.F.Monitoring of coastal dunes and lagoons: Important ecosystems to safeguard. Environments 2023, 10, 211. 10.3390/environments10120211

9. Aheto, D.W.; Mensah, E.; Aggrey-Fynn, J.; Obodai, E.A.; Mensah, C.J.; Okyere, I.; Aheto, S.P.K. Spa-tio-temporal analysis of two coastal wetland systems in Ghana: Addressing ecosystem vulnerability and implications for fisheries development in the context of climate and land use changes. Arch. Appl. Sci. Res. 2011, 3, 499–513.

10. Biney, C.A. Preliminary survey of the state of pollution of the coastal environment of Ghana. Oceanol. Acta 1982, 8, 39–43.

11. Karikari, A.; Asante, K.; Biney, C. Water quality characteristics at the estuary of Korle Lagoon in Ghana. West Afr. J. Appl. Ecol. 2009, 10. 10.4314/wajae.v10i1.45700

12. Mattah, P.A.D.; Akwetey, M.F.A.; Abrokwah, S.; Prah, P.; Tuffour, D.K.; Aheto, D.W.; Subramanian, S. Perspectives on drivers of biodiversity and environmental changes in the Keta Lagoon Ramsar Site of Ghana. Sustainability 2024, 16, 666. 10.3390/su16020666

13. Armah, A.F.; Luginaah, I.; Essandoh, P.K.; Afrifa, E.K.A. Ecological health status of the Fosu Lagoon, Southern Ghana I: Biotic assessment. J. Ecosyst. Ecogr. 2012, 2, 110–119. 10.4172/2157-7625.1000110

14. Doamekpor, L.K.; Abusa, Y.; Ketemepi, H.K.; Klake, R.K.; Doamekpor, M.E.A.M.; Anom, P.A.; Obeng, J. Assessment of heavy metals in water and sediments of Sakumo II, Chemu and Kpeshie Lagoons - Ghana. West Afr. J. Appl. Ecol. 2018, 26, 56–71. 10.4314/wajae.v26i2

15. Nartey, V.K.; Edor, K.A.; Doamekpor, L.K.; Bobobee, L.H. Nutrient load of the Sakumo Lagoon at the Sakumo Ramsar Site in Tema, Ghana. West Afr. J. Appl. Ecol. 2012, 19, 93–105.

16. Nixon, S.W.; Buckley, B.A.; Granger, S.L.; Entsua-Mensah, M.; Ansa-Asare, O.; White, M.J.; Mensah, E. Anthropogenic enrichment and nutrients in some tropical lagoons of Ghana, West Africa. Ecol. Appl. 2007, 17, 144–164. 10.1890/05-0684.1

17. Agbemehia, K. Effects of industrial waste effluents discharged into Sakumo II Lagoon in Accra, Ghana. Master’s Thesis, Kwame Nkrumah University of Science and Technology, Accra, Ghana, 2014.

18. Tay, C.K.; Asmah, R.; Biney, C.A. Trace metal levels in water and sediment from the Sakumo II and Muni Lagoons, Ghana. West Afr. J. Appl. Ecol. 2009, 16, 75–94. 10.4314/wajae.v16i1.55870

19. Pauly, D. On the ecology of a small West-African lagoon. Ber. Dtsch. Wiss. Komm. Meeresforsch. 1975, 24, 46–62.

20. Ramsar Convention Secretariat. Ramsar Information Sheet: Sakumo Ramsar Site (Site No. 565, Ghana). 2024. Available online: https://rsis.ramsar.org/RISapp/files/RISrep/GH565RIS_2405_en.pdf (accessed on 1 April 2026).

21. Ntiamoa-Baidu, Y.; Gordon, C. Coastal Wetlands Management Plans: Ghana; World Bank and Environmental Protection Council: Accra, Ghana, 1991.

22. Ramsar Convention Secretariat. Convention on Wetlands of International Importance Especially as Waterfowl Habitat (Ramsar Convention). 1971. Available online: https://www.ramsar.org/sites/default/files/documents/library/current_convention_text_e.pdf (accessed on 30 March 2026).

23. Webb, J.E. The ecology of Lagos Lagoon. I. The lagoons of the Guinea Coast. Philos. Trans. R. Soc. Lond. B. Biol. Sci. 1958, 241, 307–317.

24. Yankson, K.; Obodai, E.A. An update of the number, types and distribution of coastal lagoons in Ghana. J. Ghana Sci. Assoc. 1999, 3, 26–31.

25. Pauly, D. The biology, fishery and potential for aquaculture of Tilapia melanotheron in a small West African lagoon. Aquaculture 1976, 7, 33–49.

26. Dieckmann, U. Fishing drives rapid evolution. Sustainability 2006, 3–4, 18–19.

27. Pauly, D.; Moreau, J.; Palomares, M.L.D. Detritus and energy consumption and conversion efficiency of Sarotherodon melanotheron in a West African lagoon. J. Appl. Ichthyol. 1988, 4, 150–153.

28. Olaosebikan, B.D.; Raji, A. Field Guide to Nigerian Freshwater Fishes; Federal College of Freshwater Fisheries Technology: New Bussa, Nigeria, 1998.

29. Pauly, D. On some features of the infestation of the mouth-brooding fish Tilapia melanotheron by the parasitic copepod Paenodes lagunaris. Beaufortia 1974, 22, 9–15.

30. Pauly, D. Gasping Fish and Panting Squids: Oxygen, Temperature and the Growth of Water-Breathing Animals, 2nd ed.; International Ecology Institute: Oldendorf/Luhe, Germany, 2019.

31. Pauly, D. Spatial modelling of trophic interactions and fisheries impacts in coastal ecosystems: A case study of Sakumo Lagoon, Ghana. In The Gulf of Guinea Large Marine Ecosystem; McGlade, J.; Cury, P.; Koranteng, K.A.; Hardman-Mountford, N.J., Eds.; Elsevier Science: Amsterdam, The Netherlands, 2002; pp. 289–296.

32. Pauly, D. Anecdotes and the shifting baseline syndrome of fisheries. Trends Ecol. Evol. 1995, 10, 430.

33. Gell, P.A.; Finlayson, C.M.; Kumar, R. Ecological Character under the Ramsar Convention on Wetlands: A Sea Anchor or a Raft of Options? Wetlands 2026, 46, 29. 10.1007/s13157-026-02048-5

34. Gordon, N.; García-Rodríguez, F.; Adams, J.B. Paleolimnology of a coastal lake on the southern Cape coast of South Africa: Sediment geochemistry and diatom distribution. Journal of African Earth Sciences 2012, 75, 14–24. 10.1016/j.jafrearsci.2012.06.008

35. Stager, J.C.; Cumming, B.F.; Meeker, L.D. A 10,000-year high-resolution diatom record from Pilkington Bay, Lake Victoria, East Africa. Quaternary Research 2003, 59, 172–181. 10.1016/S0033-5894(03)00008-5

36. Liu, B.; McLean, C.E.; Long, D.T.; Steinman, A.D.; Stevenson, R.J. Eutrophication and recovery of a lake inferred from sedimentary diatoms originating from different habitats. Science of the Total Environment 2018, 628–629, 1352–1361. 10.1016/j.scitotenv.2018.02.174

37. Manjarrez-Rangel, C.S.; Halac, S.R.; Mengo, L.D.V.; Piovano, E.L.; Zanor, G.A. Paleolimnological approaches to track anthropogenic eutrophication in lacustrine systems across the American continent: A review. Limnological Review 2025, 25, 1–22. 10.3390/LIMNOLREV25030033

38. Agyepong, G.T. Coastal Wetland Management Project—Management Plan for Sakumo Ramsar Site; Wildlife Department: Accra, Ghana, 1999.

39. Agbemehia, K.; Fei-Baffoe, B. Effects of industrial waste effluents on the quality of Sakumo II Lagoon in Accra, Ghana. In Proceedings of the Conference on Fisheries and Coastal Environment; Book of Abstracts; 2017; p. 33.

40. Ansa-Asare, O.D.O.; Mensah, E.; Entsua-Mensah, M.; Biney, C.A. Impact of human activity on nutrient and trophic status of some selected lagoons in Ghana. West Afr. J. Appl. Ecol. 2008, 12, 1–10. 10.4314/wajae.v12i1.45761

41. Asmah, R.; Dankwa, H.; Biney, C.A.; Amankwah, C.C. Trends analysis relating to pollution in Sakumo Lagoon, Ghana. Afr. J. Aquat. Sci. 2008, 33, 87–93. 10.2989/AJAS.2007.33.1.11.395

42. Biney, C.A. A review of some characteristics of freshwater and coastal ecosystems in Ghana. Hydrobiologia 1990, 208, 45–53. 10.1007/BF00008442

43. Nonterah, C.; Xu, Y.; Osae, S.; Akiti, T.T.; Dampare, S.B. A review of the ecohydrology of the Sakumo wetland in Ghana. Environ. Monit. Assess. 2015, 187, 671. 10.1007/s10661-015-4872-0

44. Ntiamoa-Baidu, Y. Conservation of coastal lagoons in Ghana: The traditional approach. Landsc. Urban Plan. 1991, 20, 41–46. 10.1016/0169-2046(91)90089-5

45. Covich, A.P.; Palmer, M.A.; Crowl, T.A. The role of benthic invertebrate species in freshwater ecosystems: Zoobenthic species influence energy flows and nutrient cycling. BioScience 1999, 49, 119–127. 10.2307/1313537

46. Tait, R.V.; Dipper, F.A. Elements of Marine Ecology, 4th ed.; Butterworth-Heinemann Ltd.: Oxford, UK, 1998.

47. Townsend, D.W. Oceanography and Marine Biology: An Introduction to Marine Science; Sinauer Associates, Inc.: Sunderland, MA, USA, 2012.

48. Koranteng, K.A. Ghana Coastal Wetlands Management Project Environmental Baseline Studies—Sakumo Ramsar Site; Fisheries Department: Accra, Ghana, 1995.

49. Yankson, K. Aspects of conchological features of Anadara senilis in relation to the nature of substratum. J. Ghana Sci. Assoc. 2000, 2, 123–128.

50. Brinkhurst, R.O. A Guide for the Identification of British Aquatic Oligochaetes, 2nd ed.; Freshwater Biological Association: Ambleside, UK, 1971.

51. Brinkhurst, R.O. Guide to the Freshwater Aquatic Microdrile Oligochaetes of North America; Can. Spec. Publ. Fish. Aquat. Sci.: Ottawa, Canada, 1986; Vol. 84; p. 259.

52. Gerber, A.; Gabriel, M.J.M. Aquatic Invertebrates of South African Rivers: Field Guide, 1st ed.; Institute of Water Quality Studies: Pretoria, South Africa, 2002.

53. Yankson, K.; Kendall, M. A Student’s Guide to the Fauna of Seashores in West Africa; Darwin Initiative: Newcastle, UK, 2001.

54. WoRMS Editorial Board. World Register of Marine Species. Available online: https://www.marinespecies.org (accessed on 20 February 2025).

55. Palomares, M.L.D.; Pauly, D., Eds. SeaLifeBase. World Wide Web electronic publication. Available online: https://www.sealifebase.org (accessed on 20 February 2025).

56. Dadzie-Paintsil, E.; Mensah, J.V. Effects of urbanization on coastal wetlands in the Sekondi-Takoradi Metropolis, Ghana. Indo Pac. J. Ocean Life 2022, 6, 94–105.

57. Akintola, S.L.; Anetekhai, M.A.; Lawson, E.O. Some physicochemical characteristics of Badagry Creek, Nigeria. West Afr. J. Appl. Ecol. 2011, 18, 95–107.

58. Das, R.; Samal, N.R.; Roy, P.K.; Mitra, D. Role of electrical conductivity as an indicator of pollution in shallow lakes. Asian J. Water Environ. Pollut. 2006, 3, 143–146.

59. Horne, A.J.; Goldman, C.R. Limnology, 2nd ed.; McGraw-Hill, Inc.: New York, NY, USA, 1994.

60. Failla, A.J.; Vasquez, A.A.; Fujimoto, M.; Ram, J.L. The ecological, economic and public health impacts of nuisance chironomids and their potential as aquatic invaders. Aquat. Invasions 2015, 10, 1–15. 10.3391/ai.2015.10.1.01

61. Broza, M.; Halpern, M.; Gahanma, L.; Inbar, M. Nuisance chironomids in wastewater stabilization ponds: Monitoring and action threshold assessment based on public complaints. J. Vector Ecol. 2003, 28, 31–36.

62. Frouz, J.; Matena, J.; Ali, A. Survival strategies of chironomids (Diptera: Chironomidae) living in temporary habitats: A review. Eur. J. Entomol. 2003, 100, 459–466. 10.14411/eje.2003.069

63. De Souza Beghelli, F.G.; dos Santos, A.C.A.; Urso-Guimarães, M.V.; do Carmo Calijuri, M. Relationship between spatial distribution of the benthic macroinvertebrate community and trophic state in a Neotropical reservoir (Itupararanga, Brazil). Biota Neotrop. 2012, 12, 114–124. 10.1590/s1676-06032012000400012

64. Wright, S.; Tidd, W.M. Summary of limnological investigations in western Lake Erie in 1929 and 1930. Trans. Am. Fish. Soc. 1933, 63, 271–285.

65. Adámek, Z.; Maršálek, B. Bioturbation of sediments by benthic macroinvertebrates and fish and its implication for pond ecosystems: A review. Aquaculture International 2013, 21, 1–17. 10.1007/s10499-012-9527-3

66. Martins, A.D.; Barros, F. Ecological functions of polychaetes along estuarine gradients. Frontiers in Marine Science 2022, 9, 780318. 10.3389/fmars.2022.780318

67. Ghana News Agency. TMA has allowed encroachment on Sakumo Lagoon. Available online: https://www.ghheadlines.com/agency/ghana-news-agency/20170402/38681123/tma-has-allowed-encroachment-on-sakumo-lagoon (accessed on 21 February 2024).

68. Ghana News Agency. Sakumo Ramsar: A sad depleting bird’s habitat. Available online: https://gna.org.gh/2021/10/sakumo-ramsar-site-a-sad-depleting-birds-habitat (accessed on 21 February 2024).

69. Daily Guide Network. Disaster looms—Rapid construction works swallow Sakumono wetlands. Available online: https://dailyguidenetwork.com/disaster-looms-rapid-construction-works-swallow-sakumono-wetlands/ (accessed on 21 February 2024).

70. GhanaWeb. Sakumono Lagoon overflows its banks—Submerges 200 houses. Available online: https://www.ghanaweb.com/GhanaHomePage/NewsArchive/Sakumono-Lagoon-Overflows-Its-Banks-Submerges-200-Houses-1032 (accessed on 21 February 2024).

71. Africa Press. Sakumo Lagoon Ramsar site threatened by encroachers. Available online: https://www.africa-press.net/ghana/all-news/sakumo-lagoon-ramsar-site-threatened-by-encroachers (accessed on 26 February 2024).

72. He, X.; Wei, H. Biodiversity conservation and ecological value of protected areas: A review of current situation and future prospects. Frontiers in Ecology and Evolution 2023, 11, 1261265. 10.3389/fevo.2023.1261265

73. Wang, D.; de Knegt, H.J.; Hof, A.R. The effectiveness of a large protected area to conserve a global endemism hotspot may vanish in the face of climate and land-use changes. Front. Ecol. Evol. 2022, 10, 984842. 10.3389/fevo.2022.984842

74. Pulido-Chadid, K.; Virtanen, E.; Geldmann, J. How effective are protected areas for reducing threats to biodiversity? A systematic review protocol. Environ. Evid. 2023, 12, 18. 10.1186/s13750-023-00311-4

75. Gaget, E.; Le Viol, I.; Pavón-Jordán, D.; Cazalis, V.; Kerbiriou, C.; Jiguet, F.; Popoff, N.; Encarnação, V.M.F.; Erciyas-Yavuz, K.; Etayeb, K.S.; Molina, B.; Petkov, N.; Uzunova, D. Assessing the effectiveness of the Ramsar Convention in preserving wintering waterbirds in the Mediterranean. Biological Conservation 2020, 243, 108485. 10.1016/j.biocon.2020.108485

76. Ibrahim, I.; Aziz, N.A. The roles of international NGOs in the conservation of biodiversity of wetlands. Procedia Soc. Behav. Sci. 2012, 42, 242–247. 10.1016/j.sbspro.2012.04.187

77. Daily Guide Africa. Dying Korle Lagoon. Available online: https://newsghana.com.gh/dying-korle-lagoon/ (accessed on 4 April 2024).

78. Joy Online. The slow death of Accra’s Korle Lagoon. Available online: https://www.myjoyonline.com/the-slow-death-of-accras-korle-lagoon/ (accessed on 4 April 2024).

79. Modern Ghana. Chemu Lagoon, an end in sight? Available online: https://www.modernghana.com/news/124957/chemu-lagoon-an-end-in-sight.html (accessed on 4 April 2024).

80. PeaceFM Online. Chemu Lagoon dying. Available online: https://www.peacefmonline.com/pages/local/news/201710/331969.php (accessed on 4 April 2024).

81. Pulse. Photos: Current state of Chemu Lagoon floating bridge sparks fear among residents. Available online: https://www.pulse.com.gh/news/local/current-state-of-chemu-lagoon-floating-bridge-sparks-fear-among-residents-photos/pcs5b7h (accessed on 4 April 2024).

82. Catsadorakis, G.; Malakou, M.; Crivelli, A.J. Multifaceted local action for the conservation of the trans-boundary Prespa Lakes Ramsar sites in the Balkans. Mar. Freshw. Res. 2021, 72, 1554–1566. 10.1071/MF21123

83. Shine, C.; de Klemm, C. Wetlands, Water and the Law: Using Law to Advance Wetland Conservation and Wise Use; International Union for Conservation of Nature: Gland, Switzerland; Cambridge, UK, 1999.

84. Atubiga, J.A.; Donkor, E. Diminishing lagoon services in the era of urbanization: A case of Muni-Pomadze Lagoon in Ghana. Journal of Social Sciences 2022, 18, 164–170. 10.3844/jssp.2022.164.170

85. Dankwa, H.R.; Shenker, J.M.; Lin, J.; Ofori-Danson, P.K.; Ntiamoa-Baidu, Y. Fisheries of two tropical lagoons in Ghana, West Africa. Fisheries Management and Ecology 2004, 11, 379–386. 10.1111/j.1365-2400.2004.00406.x

86. Duku, E.; Mattah, P.A.D.; Angnuureng, D.B. Assessment of land use/land cover change and morphometric parameters in the Keta Lagoon Complex Ramsar Site, Ghana. Water 2021, 13, 2537. 10.3390/w13182537

87. Ekumah, B.; Armah, F.A.; Afrifa, E.K.A.; Aheto, D.W.; Odoi, J.O.; Afitiri, A.R. Geospatial assessment of ecosystem health of coastal urban wetlands in Ghana. Ocean and Coastal Management 2020, 193, 105226. 10.1016/j.ocecoaman.2020.105226

88. Ramsar Site Information Service https://rsis.ramsar.org/?f%5B0%5D=regionCountry_fr_ss%3AGabon&f%5B2%5D=montreuxListed_b%3A1&language=fr

89. GhanaWeb. Dredge the Sakumo Lagoon to save Tema. Available online: https://www.ghanaweb.com/GhanaHomePage/NewsArchive/Dredge-the-Sakumo-lagoon-to-save-Tema-446966 (accessed on 21 February 2024).

90. News Ghana. Dredging the Sakumo Lagoon will save Tema. Available online: https://newsghana.com.gh/dredging-the-sakumo-lagoon-will-save-tema/ (accessed on 21 February 2024).

91. Modern Ghana. Dredge the Sakumo Lagoon to save Tema. Available online: https://www.modernghana.com/news/697615/dredge-the-sakumo-lagoon-to-save-tema.html (accessed on 21 February 2024).

92. Citi Newsroom. Gov’t suspends demolition of unlawful structures at Sakumo Ramsar Site. Available online: https://www.citinewsroom.com/2022/11/govt-suspends-demolition-of-unlawful-structures-at-sakumo-r amsar-site/ (accessed on 1 May 2026).

93. Citi Newsroom. Sakumo Ramsar demolition stalls over fuel, equipment issues. Available online: https://citinewsroom.com/2025/05/sakumono-ramsar-demolition-stalls-over-fuel-equipment-issues/ (accessed on 1 May 2026).

94. Boadi, K.O.; Kuitunen, M. Urban waste pollution in the Korle Lagoon, Accra, Ghana. Environmental. 2002, 22, 301–309. 10.1023/A

95. Acheampong, S.M.; Ocloo, A.; Wutor, C.V.; Adamafio, N.A. Physico-chemical characteristics of water samples from selected water bodies in and around Accra, Ghana. Pollut. Res. 2014, 33, 835–841.

